# The role of cytochrome c in mitochondrial metabolism of human oocytes

**DOI:** 10.1101/2024.10.01.616010

**Authors:** Jakub Maciej Surmacki, Halina Abramczyk, Bogna Sobkiewicz, Renata Walczak-Jędrzejowska, Jolanta Słowikowska-Hilczer, Katarzyna Marchlewska

## Abstract

We investigated the biochemical composition of specific organelles in human oocyte cells at various maturation stages (GV, immature MI, MII with the first polar body, MII with giant polar body, and vacuoles) using Raman imaging. The structures analyzed included the nucleus, zona pellucida, perivitelline space, polar body, mitochondria, and cytoplasm. Raman imaging combined with chemometric classification via Cluster Analysis facilitated a comprehensive biochemical analysis of the proteomic, lipidomic, and DNA profiles of the human oocyte. Our findings indicate that cytochrome *c* is a crucial protein in oocyte mitochondria, essential for both cellular respiration (via oxidative phosphorylation) and apoptosis. Effective oocyte cell functioning requires cytochrome *c* in its redox-balanced forms: reduced and oxidized. The oxidized form of cytochrome *c* plays a pivotal role in the electron transport chain, enabling proper electron shuttling between complex III, cytochrome *c*, and complex IV. This process ensures controlled oxidative phosphorylation and ATP production, which are vital for most steps in oocyte maturation.

**Background:** In spite of progress in the identification of the fertile/infertile human oocytes in the recent decades, our understanding of molecular mechanisms occurring in the female reproductive cells did not make significant progress.

**Objective:** The present study aimed to develop our knowledge on mitochondrial metabolism in oocyte cells.

**Results:** Biochemical human oocyte analysis of proteomic-lipidomic-DNA profile by using Raman imaging combined with chemometric classification method of Cluster Analysis has been obtained. Our results show that cytochrome *c* is a key protein in oocyte mitochondria that is needed to maintain life (respiration via oxidative phosphorylation) and cell death (apoptosis).

**Materials and Methods:** In this experimental study, the nine fresh human native live oocytes from six donors collected in Salve Medica, Lodz, Poland have been investigated. The fingerprint and high-frequency region were evaluated by using Raman spectroscopy and imaging from 400 to 3500 cm^-1^. The principal component analysis method was used to visualize the difference in the Raman spectra of oocytes at different stage of maturation. The study was approved by the Bioethical Committee of the Medical University in Lodz, Poland (No RNN/83/23/KE). Written informed consent was obtained from all patients who voluntarily participated in the study. All procedures were conducted in accordance with the guidelines of the Declaration of Helsinki (2013).

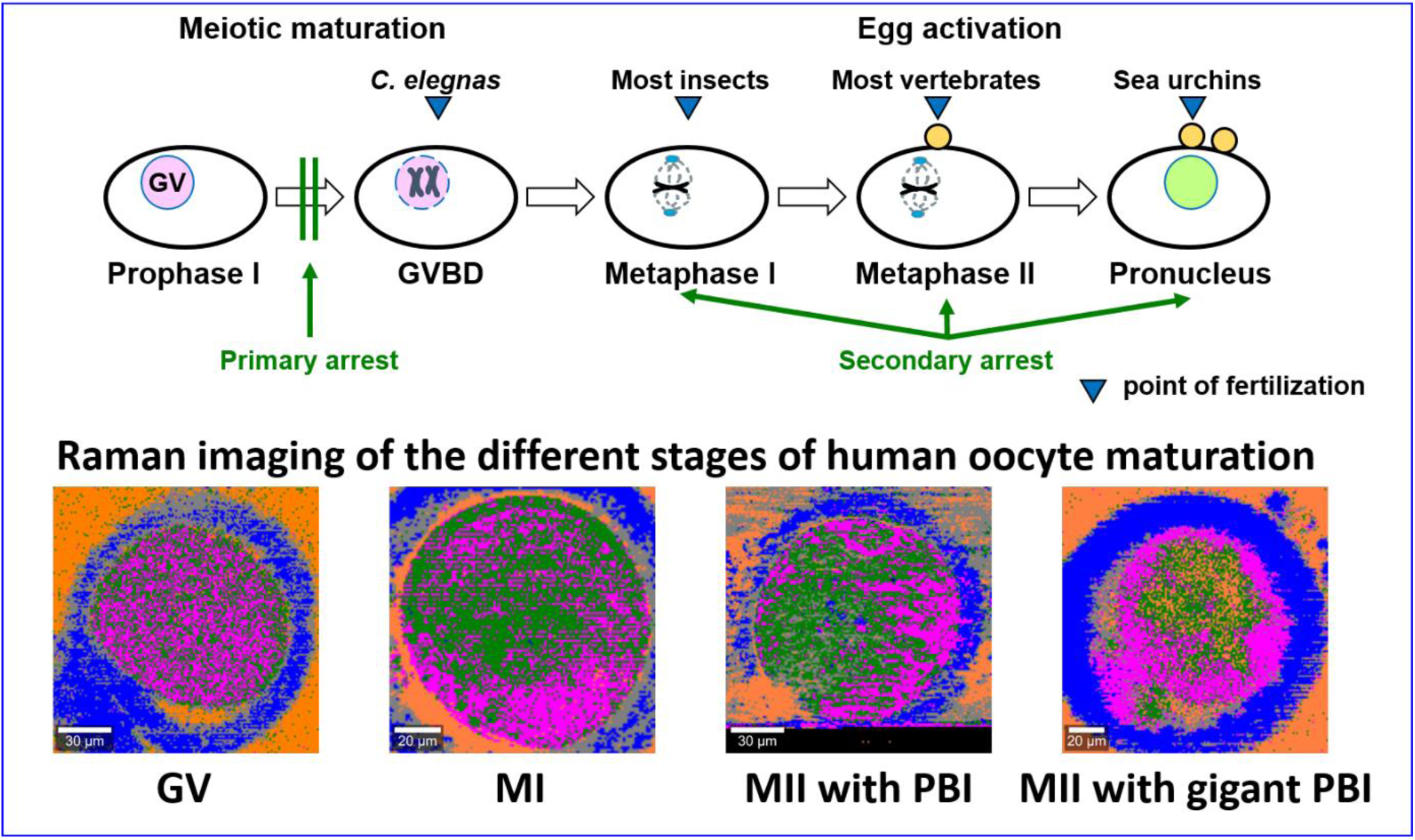

## Introduction

The formation of new life occurs through fusion of male and female gametes. This process is preceded by many mechanisms from oogenesis and spermatogenesis up to natural insemination, artificial insemination or *in vitro* fertilization. The effectiveness of insemination/fertilization depends on many dynamic and biochemical factors in both gametes. The gametes undergo many physiological changes, but prerequisite for their participation in insemination/fertilization is their own quality before starting fusion. In our previous paper we discussed the quality of male gametes in sperm. The results show that biochemical changes in sperm cells were concerned with the processes occurring in the electron transport chain in mitochondria governing the respiratory metabolism of the cells.^1^

In this paper we will focus on quality of female gametes - oocytes. The normal oocytes undergo a few cycles to reach maturation and competence for fertilization by a spermatozoon. The maturation is reached just prior to ovulation. Embryologists determine three stages to assess oocyte maturity: germinal vesicle (GV), metaphase 1 (MI) and metaphase 2 (MII) corresponding to meiosis, in which the oocyte extrudes a polar body and is ready for fertilization or cryopreservation. Once meiosis gets started, the oocytes undergo five stages called the leptotene, zygotene, pachytene stages, diplotene, and diakinesis.^2–4^

Each stage is associated with more or less dramatic alterations in bio-composition, the oocyte chromatin structure and lipid profile. The rearrangements of chromatin depend on epigenetic modifications such as histone acetylation, phosphorylation, methylation, glycosylation, ubiquitination, and sumoylation which decide about the quality of oocyte maturation.^2^

During each stage (GV, MI, MII) of meiosis, various abnormalities can occur, governed by biochemical changes in the chemical composition of the oocytes, epigenetic modifications, and dynamic factors. Oocyte maturation abnormalities are classified into four types: Type I (GV arrest), Type II (MI arrest), Type III (MII arrest), and Type IV (Mixed arrest). Uncovering the regulatory mechanisms of reproduction will enhance our understanding of the foundations of fertility and infertility.^5,6^

The most popular markers of assessing oocytes fertilization competence are based on some morphological features observed with the light microscopy.^7^ This method has many limitations because it does not provide any information on biochemistry of the process. Therefore, a variety of other methods have been proposed to examine quality of female gametes such as electron and confocal fluorescence microscopies^7,8^ as well as new omic technologies of molecular biology such as proteomics, transcriptomics, metabolomics, lipidomics.^9^ Although they provide many structural and molecular information, these methodologies have also many limitations. Undoubtedly, the most important bottleneck is the lack of information regarding biochemical composition of specific organelles in oocytes due to manipulation and processing such as invasive labeling, cellular fixation and other isolation protocols that lead to destruction of the oocyte.

Mitochondria are cytoplasmic organelles with specialized functions that play a crucial role in many processes such as energy production, calcium regulation, and apoptosis. In oocytes, the entire population of mitochondria is inherited maternally, meaning their dysfunction cannot be compensated for by male mitochondria after fertilization. Therefore, disruption of mitochondrial quality control mechanisms will lead to reproductive failure.^10^ Given their key role in the development of oocytes and embryos, research aimed at finding markers of mitochondrial function appears to be very important and timely.

Raman spectroscopy and imaging is an interesting alternative for the currently existing methods, which allows for a non-invasive monitoring of cellular processes. Raman spectroscopy and imaging has many advantages over conventional biology which needs cell disruption and release the cellular structures. In Raman imaging we do not need to destroy cells to learn about their biochemical composition and localization. Therefore, vibrational spectroscopy (IR, Raman) have been successfully applied to analyze the female gamete in animal models and in human samples.^11–18,18,19^ Unfortunately, in spite of progress in the identification of the fertile/infertile oocytes in the recent decades, our understanding of molecular mechanisms occurring in the female gametes did not make significant progress.

In this paper we will apply Raman imaging to investigate alterations of chemical composition, distribution of biochemical components and evolution of molecular processes related to oocyte growing at different stages of maturation. We will focus on the role of cytochrome *c* in the electron transport chain in mitochondria governing the respiratory metabolism of oocytes at different stages of maturation (GV, MI, MII). The paper aims to answer the question of the mechanism(s) governing electron transport chain and oxidative phosphorylation by studying alterations in electronic and vibrational spectra of cytochrome *c* in oocytes at different stages of maturation.

## Results

Scheme 1 shows the main components such as corona radiata, zona pellucida and perivitelline space, cortical granules, polar body, nucleus in meiotic metaphase II cytoplasm of the oocyte structure.

**Scheme 1.**
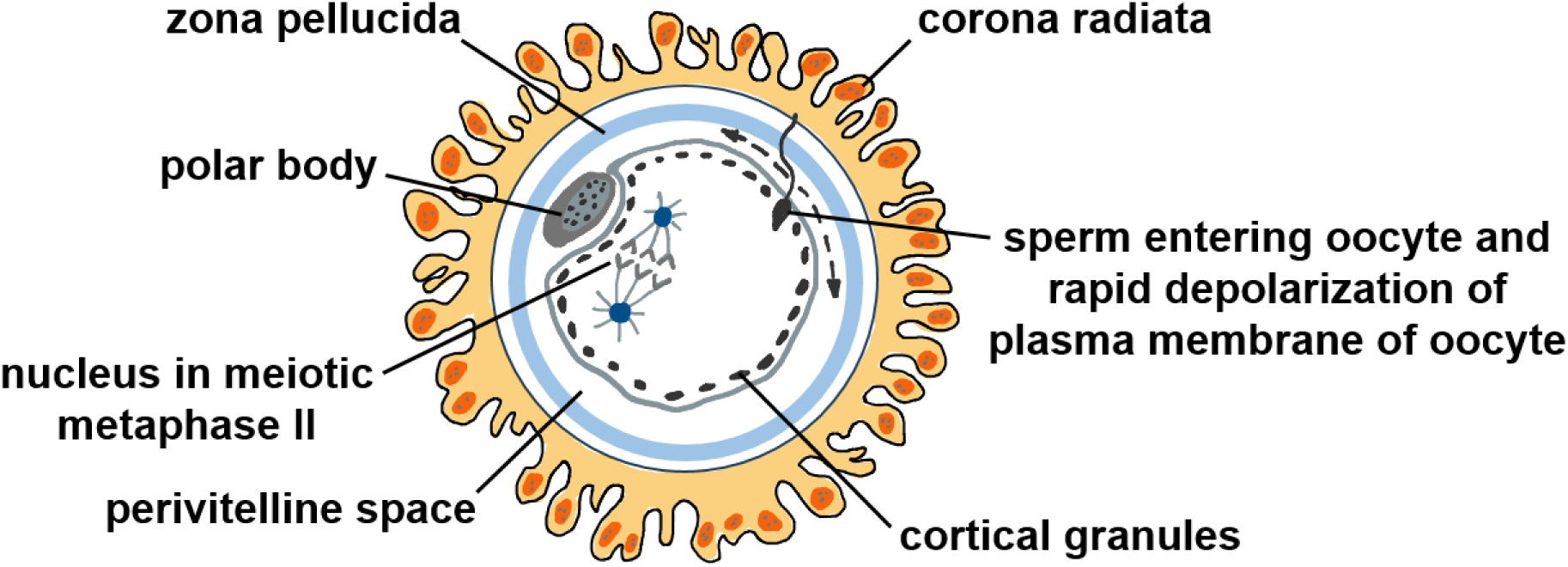
The main components such as corona radiata, zona pellucida and perivitelline space, cortical granules, polar body, nucleus in meiotic metaphase II cytoplasm of the oocyte structure.

Figure 1 shows a microscopy image, Raman image and Raman spectra of a typical immature oocyte in GV stage following the removal of the cumulus –corona cells. Germinal vesicle GV is an immature oocyte at the embryonic vesicle stage indicating the arrest of meiosis (cell division) at the prophase I stage. GV oocytes are not used for the in vitro procedures and are rejected.^20,21^

**Figure 1.**
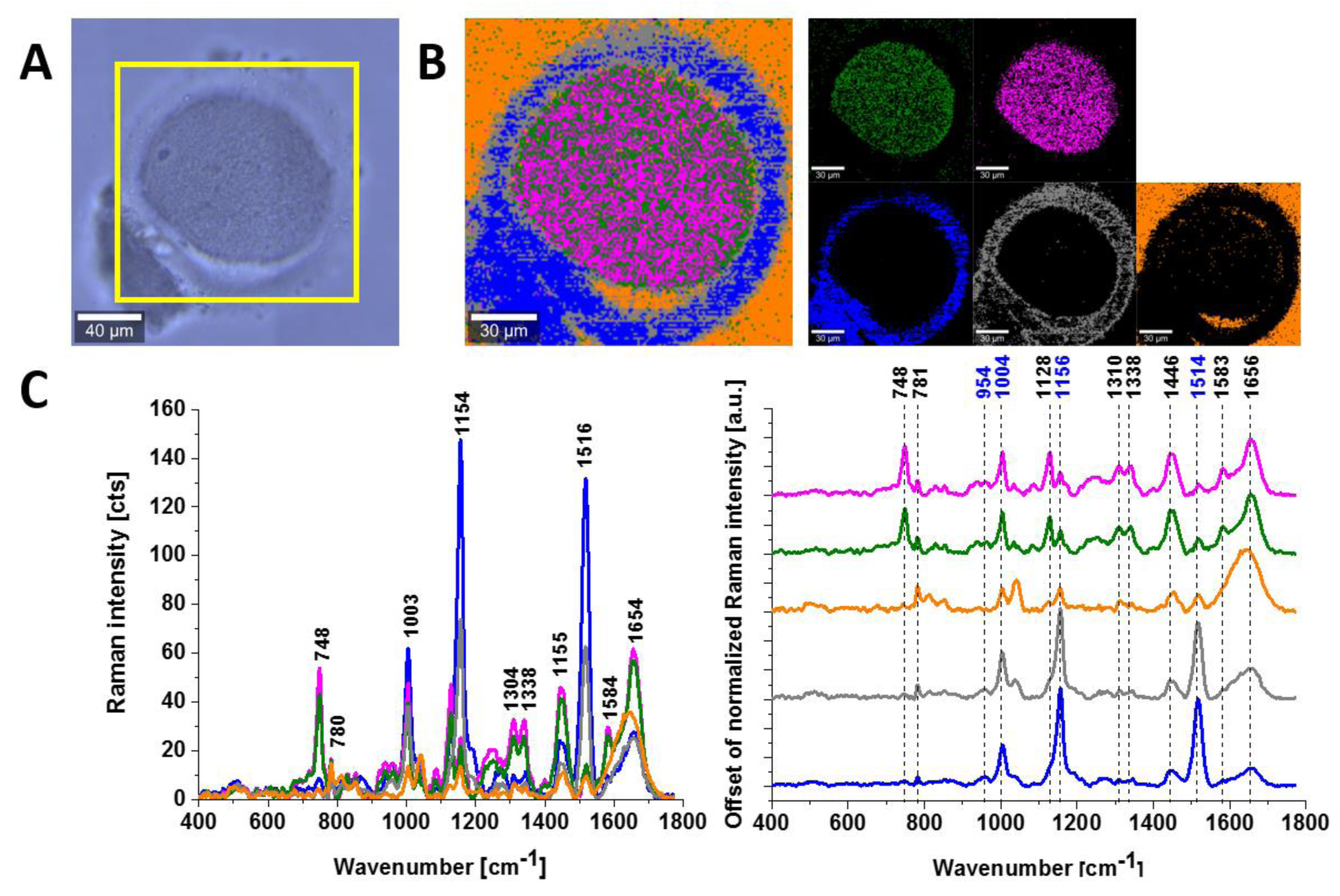
Immature oocyte in GV stage (germinal vesicle GV) following the removal of the cumulus – corona cells. (A) microscopy image, (B) Raman image obtained from the Cluster Analysis and the separate cluster components, (C) Raman spectra. The colors of spectra correspond to the colors of classes in the Raman maps. Resolution of Raman images is 1 μm, integration time 0.3 s, 10 mW at 532 nm.

The enzyme used for denudation in in vitro protocols is hyaluronidase, which removes the cumulus–corona cells (Scheme 1) as this enzyme digests the hyaluronic acid interspaced between the cumulus cells.^22^

Detailed inspection of Raman image of an immature oocyte in GV stage in Fig. 1B show a few layers of the oocyte that has been identified by Raman spectroscopy such as zona pellucida (blue colour) and perivitelline space (orange colour inside a cell).^21^ The perivitelline space is between the zona pellucida and the oocyte membrane. The green colour inside the cell represents mitochondria, the magenta represents cytoplasm. The orange colour outside the cell comes external matrix, particularly from hyaluronidase used for denudation in in vitro protocols. The nucleus of the cell in GV stage characteristic of prophase I of the first meiotic division is not visible both in the microscopy and Raman images.

Figure 2 shows a microscopy image, Raman image and Raman spectra of a typical denuded MI immature oocyte following the removal of the cumulus – corona cells. MI oocyte has no visible nucleus and has not as yet extruded the first polar body (PBI) and represent immature cell.^20^

**Figure 2.**
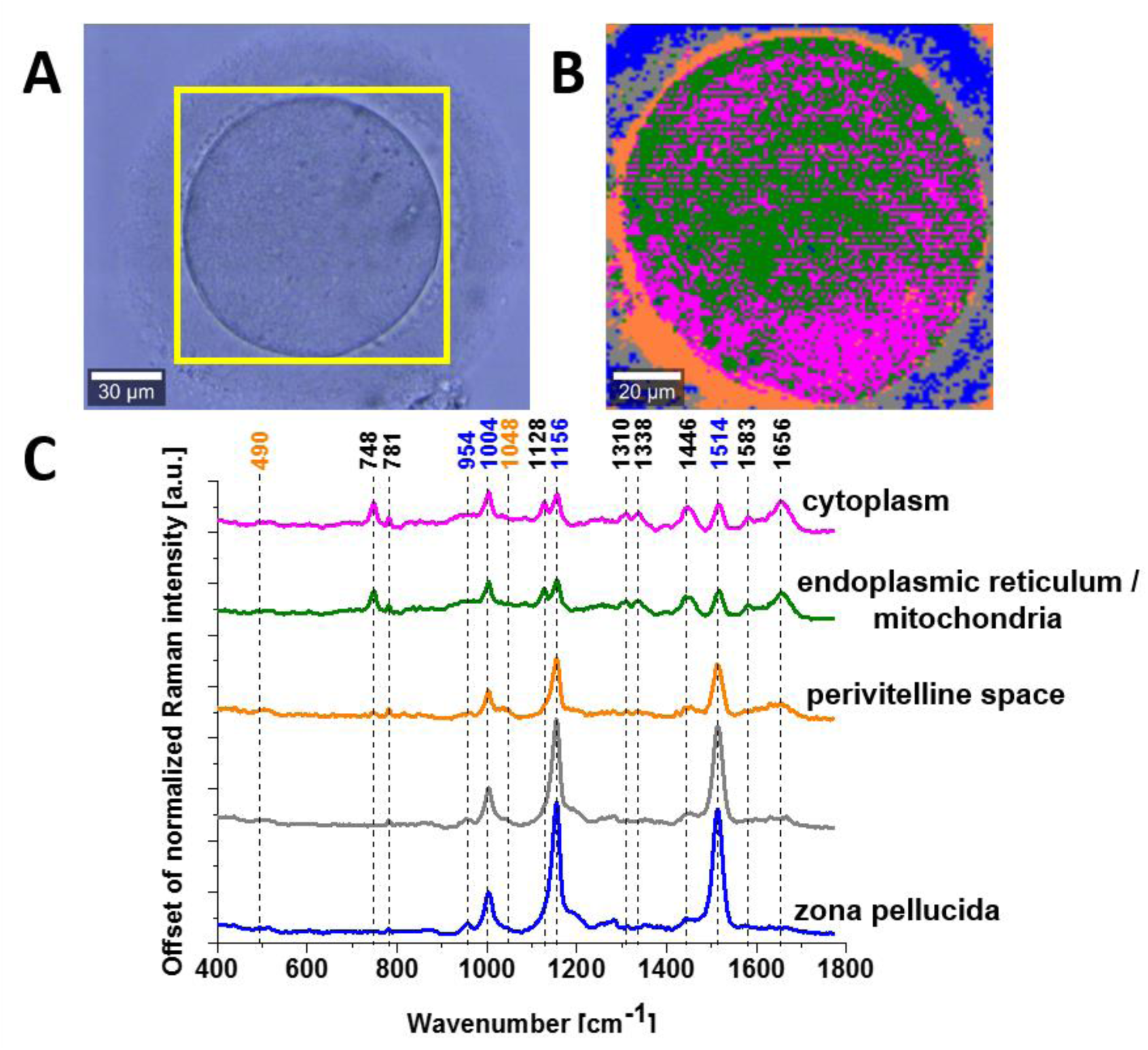
Denuded MI oocyte following the removal of the cumulus –corona radiata cells (A) microscopy image, (B) Raman image obtained from the Cluster Analysis, (C) Raman spectra. The colors of spectra correspond to the colors of classes in the Raman maps. Resolution of Raman images is 1 μm, integration time 0.3 s, 10 mW at 532 nm.

Raman images in Figures 1 and 2 combined with the vibrational features contained in the Raman spectra provides information on biocomposition of the GV and MI oocytes. The Raman spectra in Figures 1 and 2 show the distribution of carotenoids, cytochrome *c*, cytochrome *b*, lipids/cholesterol, proteins and DNA/RNA.

Detailed inspection of a typical denuded MI oocyte following the removal of the cumulus – corona cells in Figure 2B show a few layers of the oocyte that has been identified as zona pellucida (blue colour) and perivitelline space (orange colour).^21^

From the clinical practice, the perivitelline space is relevant because the cortical granules released from the cytoplasm are deposited in the perivitelline space to block polyspermy.^23^

One can see from Figure 2B that the perivitelline space layer is thick and the fluid from the cortical granules does not leak outside the perivitelline space into zona pellucida. For MI oocyte only small amount of fluid from cortical granules is released to the perivitelline space layer, because the Raman signal at 1048 cm^-1^ in the perivitelline space corresponding to polysaccharides (main component of the cortical granules) is weak.^24^ The zona pellucida both for GV and MI stages is dominated by carotenoids characterized by the bands at 1514, 1156 and 954 cm^-1^.^24^ More detailed information on these components is presented in Figures 3 and 4 for GV and MI oocytes, respectively.

**Figure 3.**
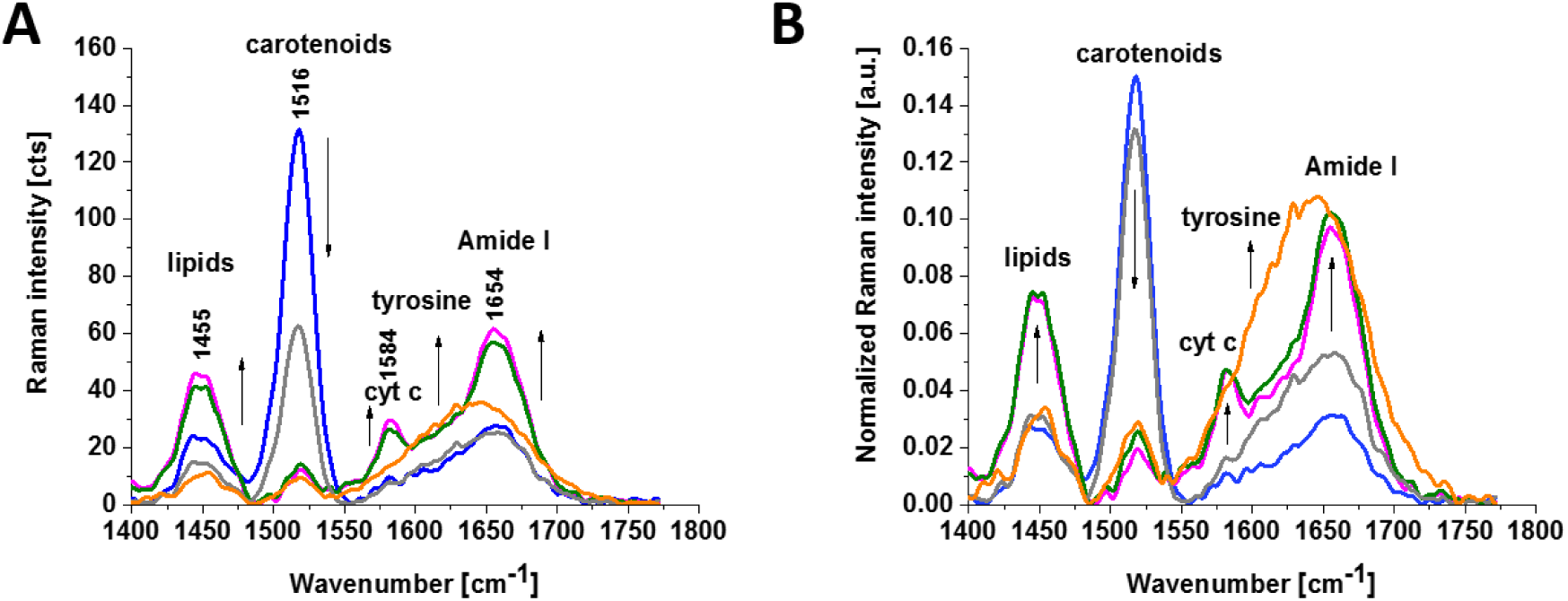
Denuded GV oocyte following the removal of the cumulus – corona radiata cells (A) Raman spectra, (B) normalized Raman spectra. The colors of spectra correspond to the colors of classes in the Raman maps in Figure 1.

**Figure 4.**
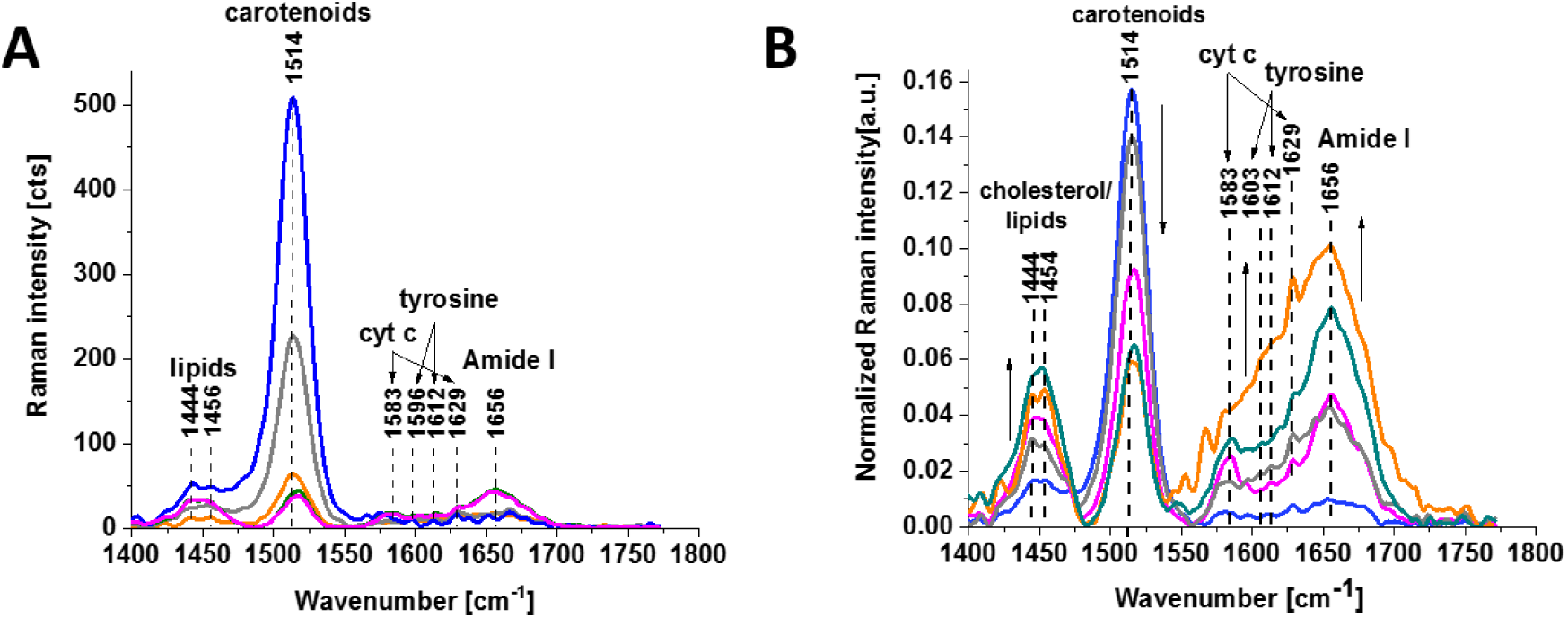
Denuded MI oocyte following the removal of the cumulus –corona radiata cells (A) Raman spectra, (B) normalized Raman spectra. The colors of spectra correspond to the colors of classes in the Raman maps in Figure 2.

One can see from Figures 2 and 6 that the amount of carotenoids decreases spectacularly in the perivitelline space, mitochondria and cytoplasm compared to the zona pellucida. In contrast, the relative amount of lipids and cholesterol (1444 and 1456 cm^-1^), cytochrome *c* (748, 1127, 1311, 1399, 1583 and 1629 cm^-1^), tyrosine (1603 and 1612 cm^-1^) and Amide I (1656 cm^-1^) increases in mitochondria and cytoplasm.^24–27^

One can see from Figures 1, 3 and 2, 4 that mitochondria of the oocytes in GV and MI stages are dominated by cytochrome *c* (748, 1127, 1311, 1399, 1583 and 1629 cm^-1^). A cytoplasm with homogenous texture and smooth appearance is dominated by lipids and cholesterol (1444 and 1456 cm^-1^) and proteins (Amide I (1656-80 cm^-1^), tyrosine (855, 1603 and 1612 cm^-1^), DNA/RNA (781, 1655-1680 cm ^-1^: T, G, C (ring breathing modes of the DNA/RNA bases).^24–27^ The oocytes contain also polysacharides (490, 540 and 1048 cm^-1^)^24^ presented in Figure 5. Some authors assigned the Raman band at 1048 cm^-1^ to PO4 symmetric stretching vibration.^14^

**Figure 5.**
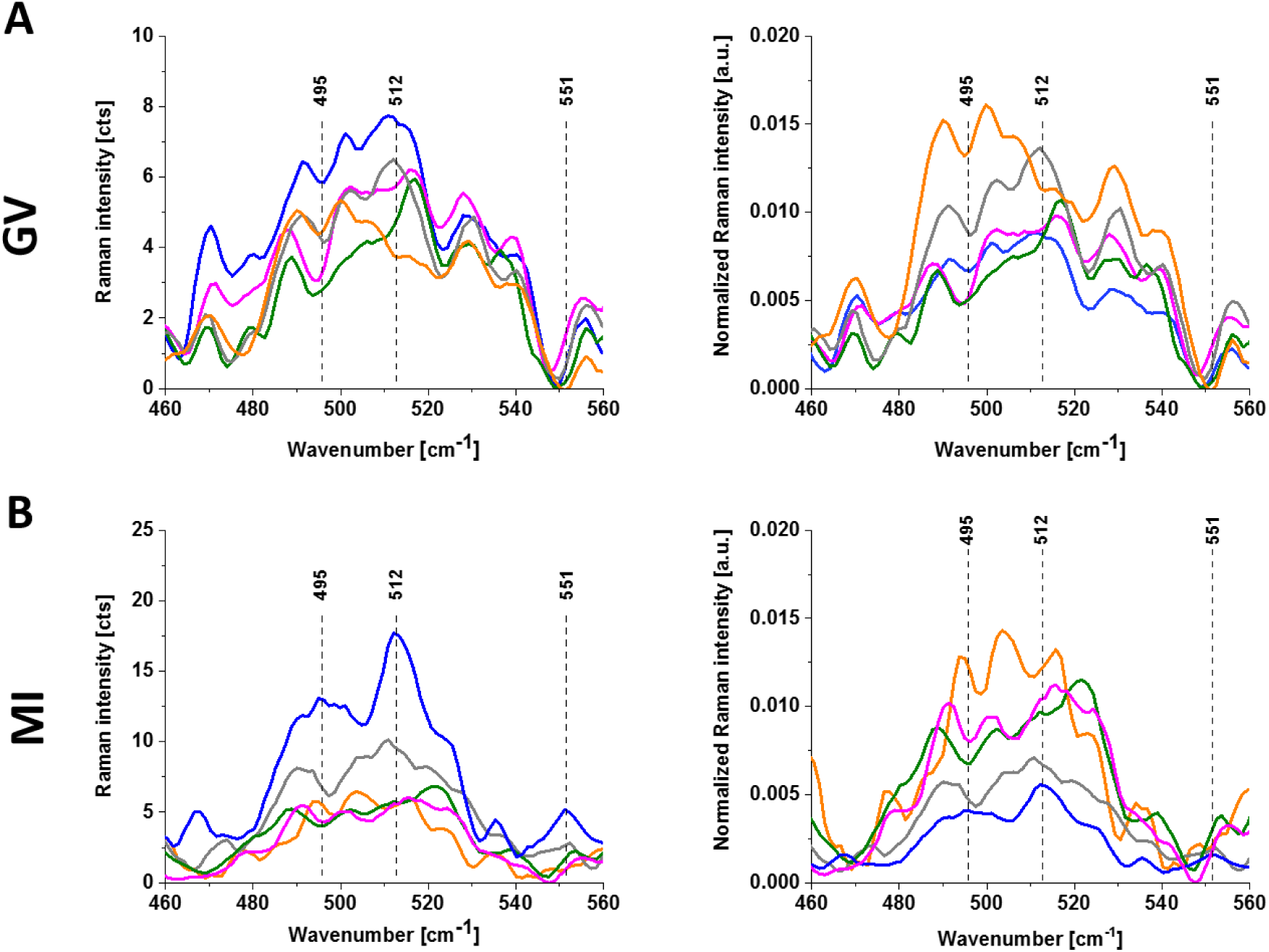
Denuded GV (A) and MI (B) oocyte non-normalized and normalized Raman spectra. The colors of spectra correspond to the colors of classes in the Raman maps in Figures 2 and 3.

Figure 6 shows a microscopy image, Raman image and Raman spectra of a typical denuded M II oocyte following the removal of the cumulus – corona cells in preparation for intracytoplasmic sperm injection (ICSI). MII oocyte has visible the first polar body (PBI) and represent a mature cell.^20,28^

**Figure 6.**
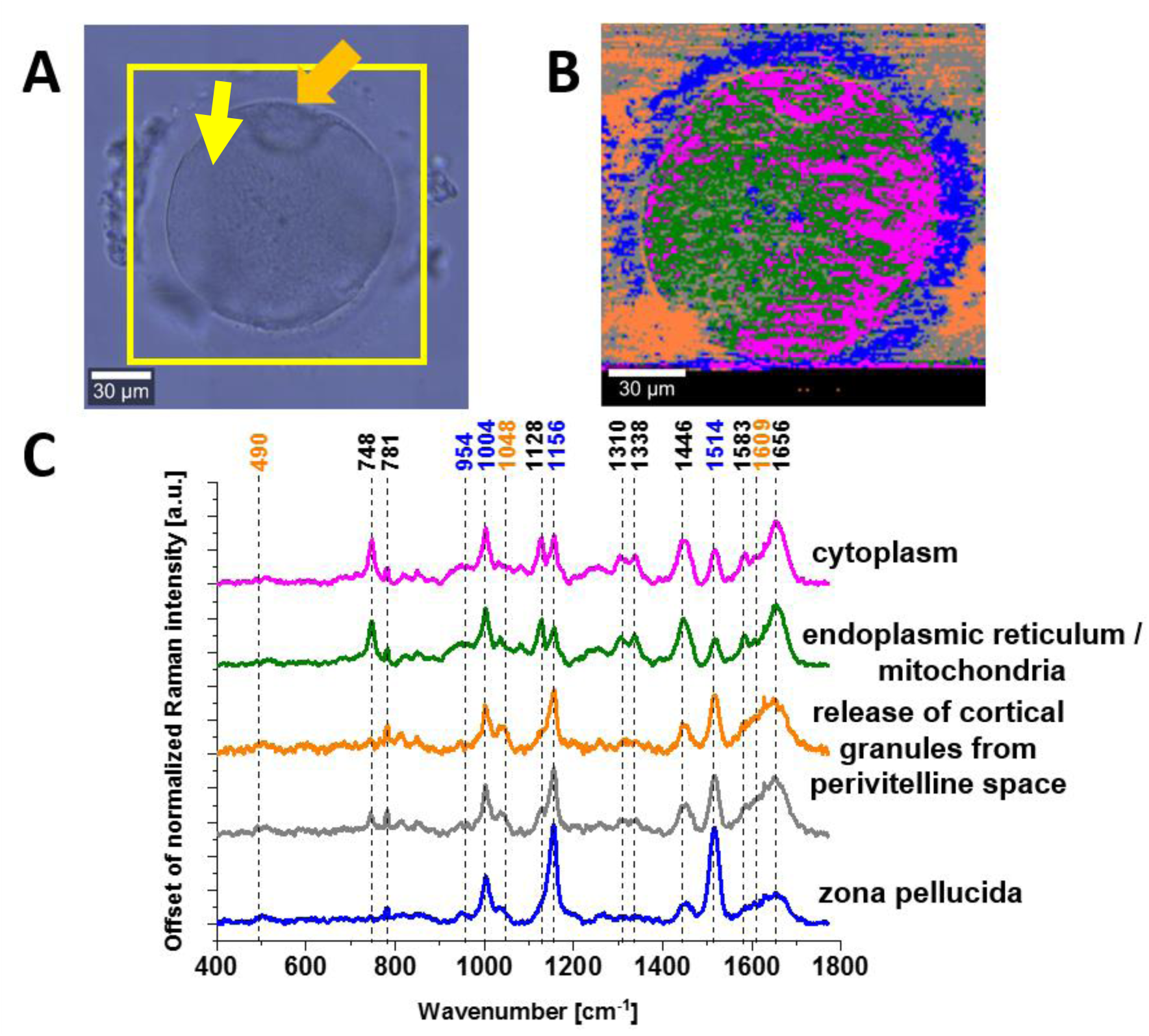
Denuded M II oocyte following the removal of the cumulus –corona radiata cells in preparation for intracytoplasmic sperm injection (ICSI) with the first normal size polar body (PB) (marked with an yellow arrow) (A) microscopy image, (B) Raman image obtained from the Cluster Analysis, (C) Raman spectra, The colors of spectra correspond to the colors of classes in the Raman maps. Resolution of Raman images is 1 μm, integration time 0.3 s, 10 mW at 532 nm.

The microscopy image and Raman image in Figure 6 show good-quality metaphase II (MII) oocyte which is characterized by a spherical shape, a single, normal-sized polar body (PB), broad zona pellucida (blue colour) and very narrow perivitelline space (orange colour inside the cell). From the clinical practice, the perivitelline space is relevant because it is the place where the polar body lodges after meiosis (yellow arrow in Figure 6).^20^

One can see from Figure 6B that for MII oocyte the fluid from the cortical granules leaks outside the perivitelline space into zona pellucida (orange colour in the 6B image). In contrast to MI immature oocyte (Figure 1), for MII mature oocyte significant amount of fluid from cortical granules is released, because the normalized Raman signal at 1048 cm^-1^ the corresponding to polysaccharides (main component of the cortical granules) is much stronger than for MI.^24^ The orange colour outside the cell comes from hyaluronidase used for denudation in in vitro protocols.

Polysaccharides released from the granules cause the space to swell, pushing the zona pellucida farther from the oocyte.^23^ The hydrolytic enzymes released by the granules cause the zona reaction, which removes the ZP3 ligands from the zona pellucida.^23^ ZP3 zona pellucida sperm-binding protein 3, also known as zona pellucida glycoprotein 3 (Zp-3) or the sperm receptor. Zp-3 induce the acrosome reaction of sperm cells at the beginning of fertilization. Zp-3 allows spermatozoa to penetrate the zona pellucida and fuse with the oocyte membrane, which is a prerequisite for fertilization of oocytes.^23^

A cytoplasm (magenta colour) has homogenous texture. The oocyte nuclear maturity is usually assessed by presence of the first polar body. However, it has been shown by using polarized light microscopy that oocytes displaying a polar body may still be immature.^20^ It was suggested that a visible meiotic spindle (MS) can be considered as mature II (MII) stage oocytes.^29^ In metaphase II (MII) oocyte the concentration of carotenoids in zona pellucida decreases significantly by factor around 5. In contrast to MI metaphase the concentration of lipids/proteins in zona pellucida becomes similar to that of the cytoplasm. The concentration of lipids, cytochrome *c* and proteins is maintained at the similar level as in the MI stage. The biocomposition of the polar body is dominated by cytochrome *c*, lipid and proteins of α helix structure.

Figure 7 shows vacuolated MII oocyte with a giant first polar body (PBI). It depicts the oocyte dysmorphisms with clear structure of cytoplasmic vacuolization. The vacuoles are organelles characteristic of eukaryotic cells and occupy a separate space in the cytoplasm, surrounded by a single protein-lipid membrane. The vacuoles remove excess water and harmful metabolic products from the cell.

**Figure 7.**
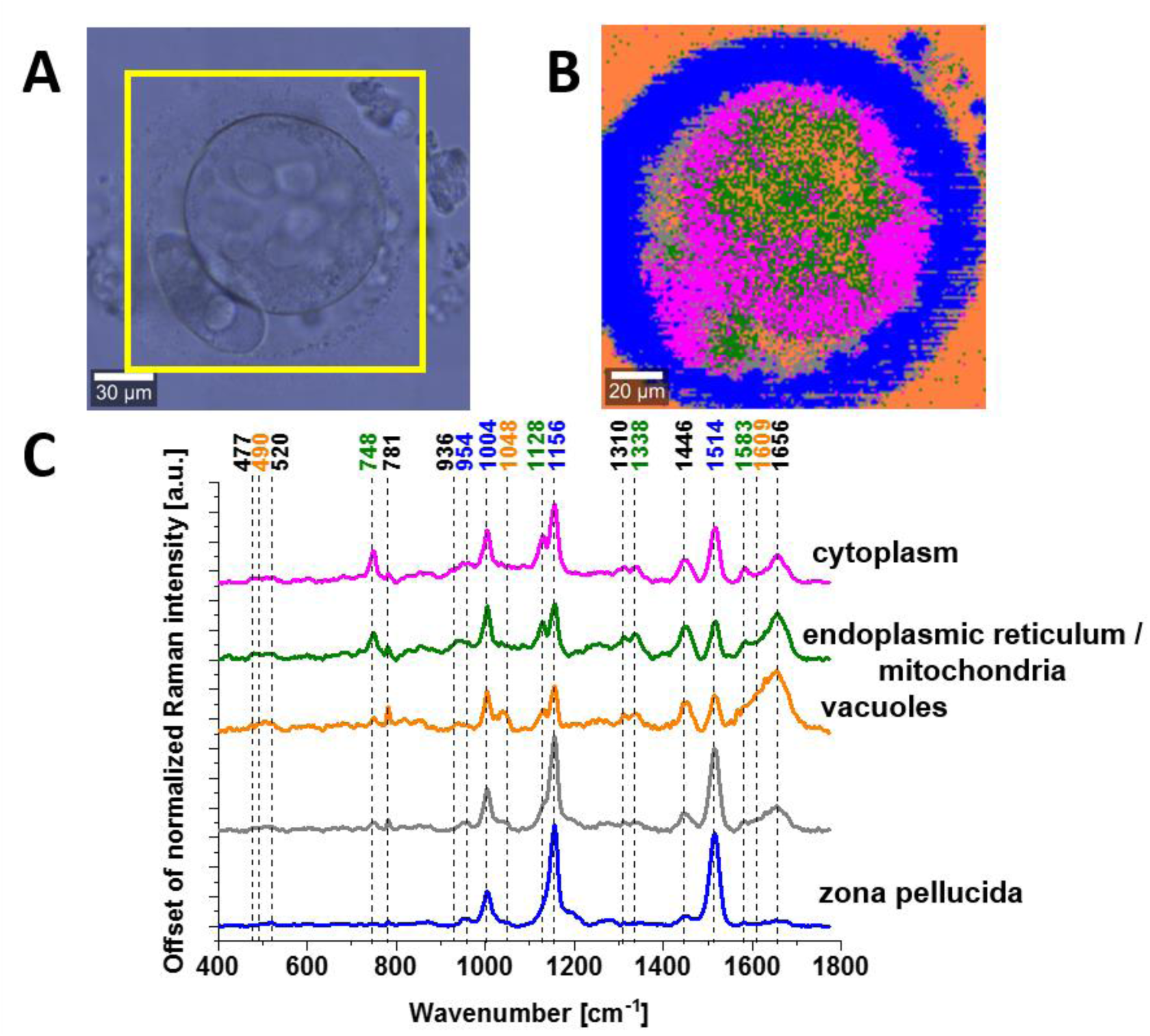
Vacuolated MII oocyte with a giant first polar body (PBI), (A) microscopy image, (B) Raman image obtained from the Cluster Analysis, (C) Raman spectra. The colors of spectra correspond to the colors of classes in the Raman maps. Resolution of Raman images is 1 μm, integration time 0.3 s, 10 mW at 532 nm.

Figure 7 shows that cytochrome *c* and *b* (green colour) is not distributed homogenously inside cytoplasm any longer as it occurs in MI metaphase (Figure 2), but it concentrates around the vacuoles.

One can see in Figure 7B Raman image of a vacuolated MII oocyte with a giant first polar body (PBI). Several small vacuoles distributed throughout the oocyte cytoplasm and the PBI (orange colour inside cytoplasm) are visible. When compared our results for the orange spectra in Figures 7 C and 2 C one can clearly state that vacuoles are filled with fluid that is virtually identical with perivitelline fluid. The orange colour outside the cell comes from hyaluronidase used for denudation in in vitro protocols.

Figure 7C shows that in contrast to other organelles the vacuoles contain large amount of polysacharides (the band of glycogen at 490 and 1048 cm^-1^) and phosphorylated proteins catalysed by tyrosine kinase (the band of phosphorylated tyrosine at 1609 cm^−1^.^30,31^ The enhanced activity of tyrosine kinase is also observed in perivitelline fluid in GV, MI and MII metaphases (Figures 1-6). Figure 7 shows that the vacuoles accumulates also proteins and lipids virtually identical with those in cytoplasm. Polysaccharides and enzymes released from the granules cause the space to swell, pushing the zona pellucida farther from the oocyte.^23^ That is why the zona pellucida in Figure 7B is thick and clearly visible. It has been reported that the hydrolytic enzymes released by the granules induces the zona reaction, triggering the acrosome reaction of sperm cells, which is a prerequisite for fertilization.^23^

## Discussion

The results presented in this study demonstrates that Raman imaging is able to explain some morphological characteristics of an ideal immature/mature human oocyte such as texture of cytoplasm, a single polar body, an appropriate zona pellucida thickness and proper perivitelline space in terms of biochemical composition. Unfortunately, in spite of progress in the identification of the fertile/infertile oocyts in the recent decades, our understanding of molecular mechanisms occurring in the female gamets did not make significant progress. Most vibrational methodologies based on Raman spectroscopy concentrated on identification and distribution of chemical components alone, which is, in fact, not enough to determine molecular mechanisms to ensure optimal conditions for subsequent fertilization.^11–18,18,19^

Figure 8 shows the proteomic-lipidomic-DNA-carotenoids profile for different stages of oocyte maturation extracted from the cytoplasm area (magenta colour in Raman images in Figures 1-7).

**Figure 8.**
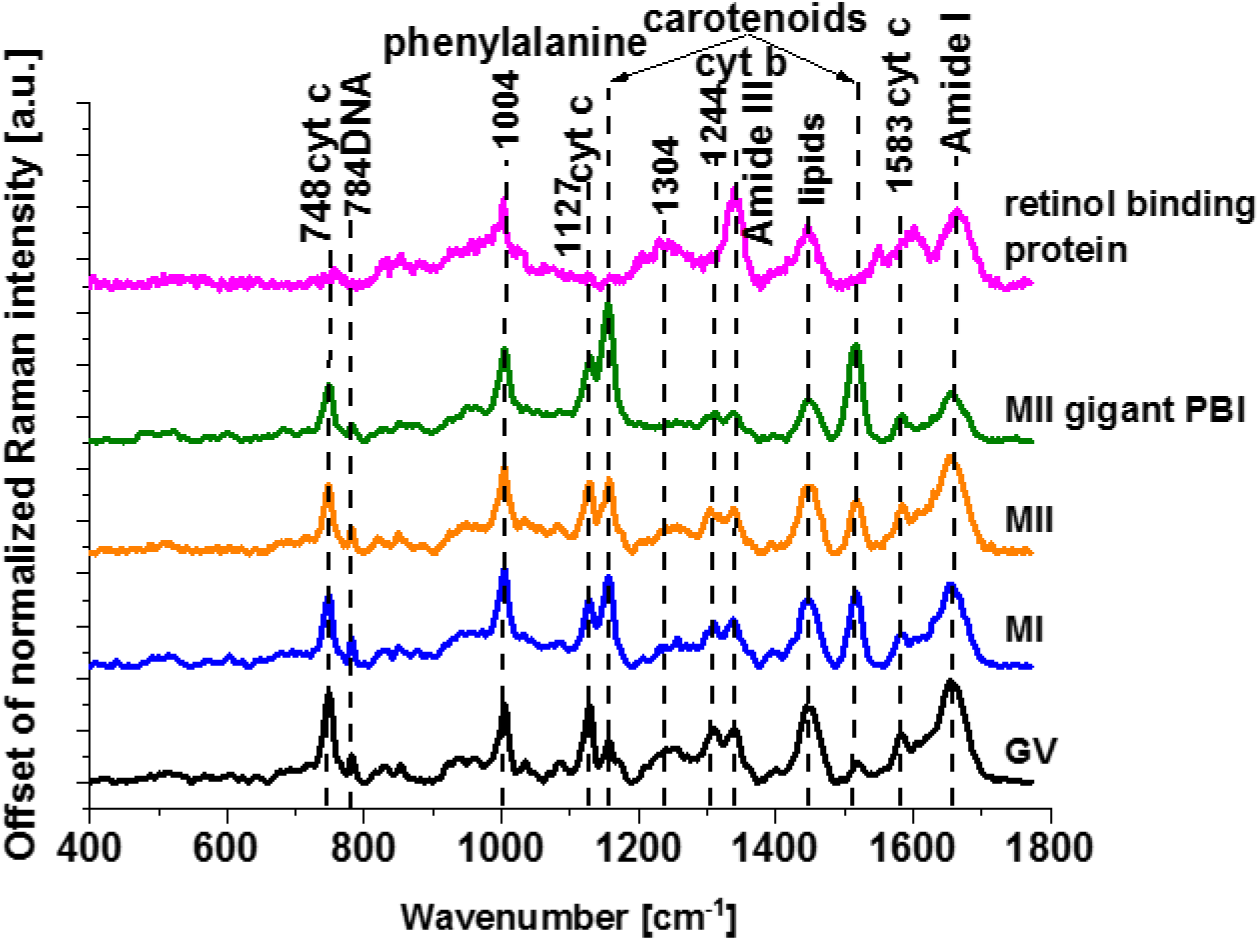
Proteomic – lipidomic – DNA - carotenoids profile for different stages of oocyte maturation in cytoplasm (magenta colour in Raman images in Figures 1-7).

Let us focus first on carotenoids. According to our best knowledge the role of carotenoids for the maturation and the development of the oocytes and cytochrome *c* in remodelling the mitochondrial electron transport chain of oocytes has not been studied yet by Raman and IR spectroscopies. Humans as other mammals are not able to synthetize carotenoids thereby they are taken from the diet. Nutrition may impact reproductive health and fertility potential. The role of dietary antioxidants on the reproductive performance and birth outcomes is a topic of emerging interest, but findings are still inconsistent. Some groups showed that vitamin C, β-carotene, and vitamin E were related to better assisted reproductive procedures techniques outcomes.^32,33^ Another group of authors found that vitamin C supplementation increased pregnancy rates primarily among non-smokers.^34^ Carotenoids serve important functions in follicular fluid as antioxidants, and regulators of steroidogenesis, folliculogenesis and oogenesis.^35,36^

In contrast, recent reports found no associations between in vitro fertilizatiion (IVF) outcomes and the intake of vitamins C, D, E, and α-carotene, β-carotene, beta-cryptoxanthin, lutein, and folate.^36^ Moreover, it has been reported that, the high intake of some vitamins (e.g. vitamin A) may result in adverse reproductive outcomes.^35^

In our measurement we use the oocytes from the IVF procedures where the applied protocols do not include carotenoids supplementation. It means that the oocytes presented in the paper contain carotenoids that are taken from the diet. In the view of our results presented so far one can state that for all stages of oocytes maturation carotenoids are present mostly in the zona pellucida. In contrast, neither carotenoids nor transformed forms of carotenoids such as retinal, retinol and retinoic acid are not observed in the perivitelline space and in cytoplasm. It may suggest that they primarily serve as antioxidants to protect oocytes for all stages of maturity from ROS processes occurring in the extracellular matrix of oocytes. How carotenoids affect mitochondrial activity and the ROS processes associated with the electron transport chain is still unknown. However, one needs to stress that vibrational spectra of carotenoids are clearly visible due to Resonance Raman enhancement effect as carotenoids have their electron absorption spectra close to the laser excitation at 532 nm used in this paper. The retinoids products of the carotenoids transformation have the electron absorption at around 350 nm from the excitation wavelength and their Raman signals are not amplified due to the Raman resonance. Thereby the used detectors may not be sensitive to detect low concentrations of retinoids inside the cytoplasm.

One can see from the Table 1 that the relative concentration of carotenoids decreases spectacularly inside the cytoplasm when compared with the zona pellucida. Table 1 shows the ratio of the Raman signal at 1516 cm^-1^ in zona pellucida (ZP) to the signal in the perivitelline space (PS), cytoplasm (C) and mitochondria (MIT). The ratio is better to study correlation between the maturity stage and the concentration of carotenoids and other components in the oocytes as the results were retrieved from different patients having different life style, diet and vitamin supplementation.

**Table 1.** The ratio of the Raman signal at 1516 cm^-1^ in zona pellucida (ZP) to the signal in the membrane (M), perivitelline space (PS), cytoplasm (C), mitochondria (MIT).

|  | ZP/M | ZP/PS | ZP/C | ZP/MIT |
| --- | --- | --- | --- | --- |
| GV | 1.1708 | 5.5700 | 7.9994 | 6.2402 |
| MI (immature) | 1.2747 | 2.3638 | 4.7853 | 5.0961 |
| MII (with PBI) | 1.7159 | 1.7659 | 2.6436 | 3.5095 |
| MII (with gigant PBI) | 1.1296 | 2.8525 | 1.7756 | 2.3836 |

Raman spectra in Figure 8 were extracted from the magenta colour in Raman images for all stages of maturation provide information on proteomic profile inside the oocyte. The proteomic profile is represented by the Amide I band (1656 cm^-1^), Amide III (1304 and 1244 cm^-1^), phenylalanine (1003 cm^-1^).^24^

We found in this paper that carotenoids are present in human oocytes, but we did not identified the products of their transformation such as retinoids. It may be due to the fact that there is no Resonance Raman signal amplification in retinoids in contrast to carotenoids. Thereby the used detectors may not be sensitive to detect low concentrations of retinoids inside the cytoplasm. This suggestion seems to have a strong support from the earlier papers, where it was reported that retinoids play a role in ovarian function.^37,38^ Retinol is transported into the oocytes by retinol-binding protein (RBP). Inside the cell, cellular retinol-binding protein (CRBP) governs retinol accumulation and metabolism.^37^ Since the actions of retinoids are mediated, in part, by retinoid-binding proteins^37^, we compared the RBP profile with proteomic profile obtained from the Raman vibrational features in Figure 8. The main band of Amide I of RBP is shifted to 1664 cm^-1^ in contrast to the bands at 1656 cm^-1^ observed in all stages of maturity of the oocytes. It indicates that the oocytes contain a significant amount of lipids that have the band at 1657 cm^-1^ (C=C) vibration) that overlap with the Amide I band.^24^

In an effort to better understand proteomic profile one can notice that RBP does not contain the band at 784 cm^-1^. The band at 784 cm^-1^ originates from DNA and represents the nucleus of the oocyte. This band is clearly visible at the first stage, where the germinal vesicle (GV) is the nucleus of the oocyte. It has been reported that GV contains chromatin (DNA) that exhibits a unique configuration regulated by dynamic histone modifications.^39–42^ Raman image in Figure 2 shows that the oocyte chromatin is located homogenously throughout the GV.

However, it was reported that the proteomic profile is more complicated and contains 12 proteins that were identified between the GV and MII stage by two-dimensional (2D) electrophoresis and mass spectrometry.^38^ It is interesting to notice that the vibration at 1583 cm^-1^ of RBP is in resonance with the vibration of cytochrome *c*. Mitochondrial activity is related to the presence of cytochrome *c*. According to our knowledge we showed for the first time distribution of cytochrome *c* in mitochondria of human oocytes. Mitochondria, which divide autonomously regardless of cell division, contain cytochrome *c*, which is located in the oocytes at each stage of maturity. The green colour in Figures 1-7 represents distribution of cytochrome *c* and *b* in mitochondria. In the spectral region between the 1583 cm^-1^ band of cytochrome *c* and RBP and the 1656-1664 cm^-1^ bands of Amide I of proteins there are bands at 1603 and 1609 cm^-1^ corresponding to tyrosine and phosphorylated tyrosine. We showed that this triangle: cytochrome *c* - tyrosine of protein kinases - RBP is related to the mitochondrial activity and respiration of cells.^1,31,43,44^

Earlier papers did not associated the band at 1603 cm^-1^ with tyrosine kinase activity, but they called it “the band of life”.^14^ Indeed, the intuition was correct because we explained how tyrosine activity is related to cytochrome *c* in the electron transport chain and oxidative phosphorylation.^1,31,43,44^ Mitochondria contain also mitochondrial DNA (mtDNA), which is inherited in humans only from the female, in contrast to nuclear genes (DNA) received from both parents. Only the female passes on her mtDNA to her offspring^45^ because only the oocyte cytoplasm, containing the mitochondria, is provided to the zygote. Sperm cells contain both genomic (nuclear) DNA and mitochondria, but mitochondria are located in the midpiece, which does not penetrate the egg. Only women who have daughters pass on mtDNA to subsequent generations (mitochondrial Eve).

Recently we showed how cancer cells remodels the mitochondrial electron transport chain through dysfunction of electron transfer between complex III - cytochrome *c* - complex IV.^1,43,44^ It has been proposed that reversible phosphorylation of cytochrome *c* mediated by cell signaling pathways are primary regulatory mechanisms that determine electron transport chain (ETC) flux, proton gradient ΔΨm, ATP production, ROS generation and mitochondrial respiration, linking oxidative phosphorylation to human cancer through a lack of energy, excess ROS production, excess cytochrome *c* release to cytoplasm, and activation of apoptosis.^1,43,44^

To understand mechanisms responsible for the redox status of cytochrom *c* let us briefly remind the role of cytochrome *c* and cytochrome *b* in the electron transfer chain (ECT) that play a key role in effectiveness of mitochondrial respiration. The cytochrome *b* is located in complex III. The cytochrome *c* is located between the complex III and the complex IV. Each cytochrome contains a heme group with an iron ion in the center, which, when receiving an electron, changes from Fe^3+^ to Fe^2+^ oxidation state. After donating the electron to the next carrier, the iron atom returns to the Fe^3+^ state. The complex III transfers electron to cytochrome *c*, which in turn transfers it to the cytochrome oxidase (complex IV) containing two cytochromes (cytochrome *a* and cytochrome *a3*) bound to two copper atoms (Cu A and Cu B), respectively. During electron transfer, the copper atoms oscillate between the Cu^2+^ state and the Cu^1+^ state.

The redox status of cytochrome *c* can be easily monitored by Raman spctroscopy analysing the vibrations of cytochrome *c*, particularly 1583 cm^-1^. We discussed this issue in details in last publications^1,43,44^. The Raman spectra from Figures 1-7 demonstrates that for all maturation stages of the oocytes, the redox balance of cytochrome *c* in the mitochondria is shifted to the Fe^3+^ oxidized state, which is indicative of an balanced electron transport chain processes. Another authors reported that mitochondrial metabolism is controlled also by suppresing complex I.^46^

Now we present principal components (PCA) analysis of all Raman data presented in Figure 9 to visualize the vibrational features for a better understanding of biochemical hallmarks of the different stages of maturation GV, MI, MII, MII giant PB of human oocyte. The distribution of the scores on principal components 1, 2, and 3 presented in Figure 9 clearly distinguishes the stages of maturation of the human oocytes cells. The loadings plots in Figure 9 helps to identify the chemical components of the oocytes. Positive PC3 values – Raman bands at 748, 1128, 1310, and 1583 cm^-1^ are characterized by an oxidized form of cytochrome *c*. Positive values on PC2 at 1516, and 1156 cm^-1^ characterize carotenoids rich region.

**Figure 9.**
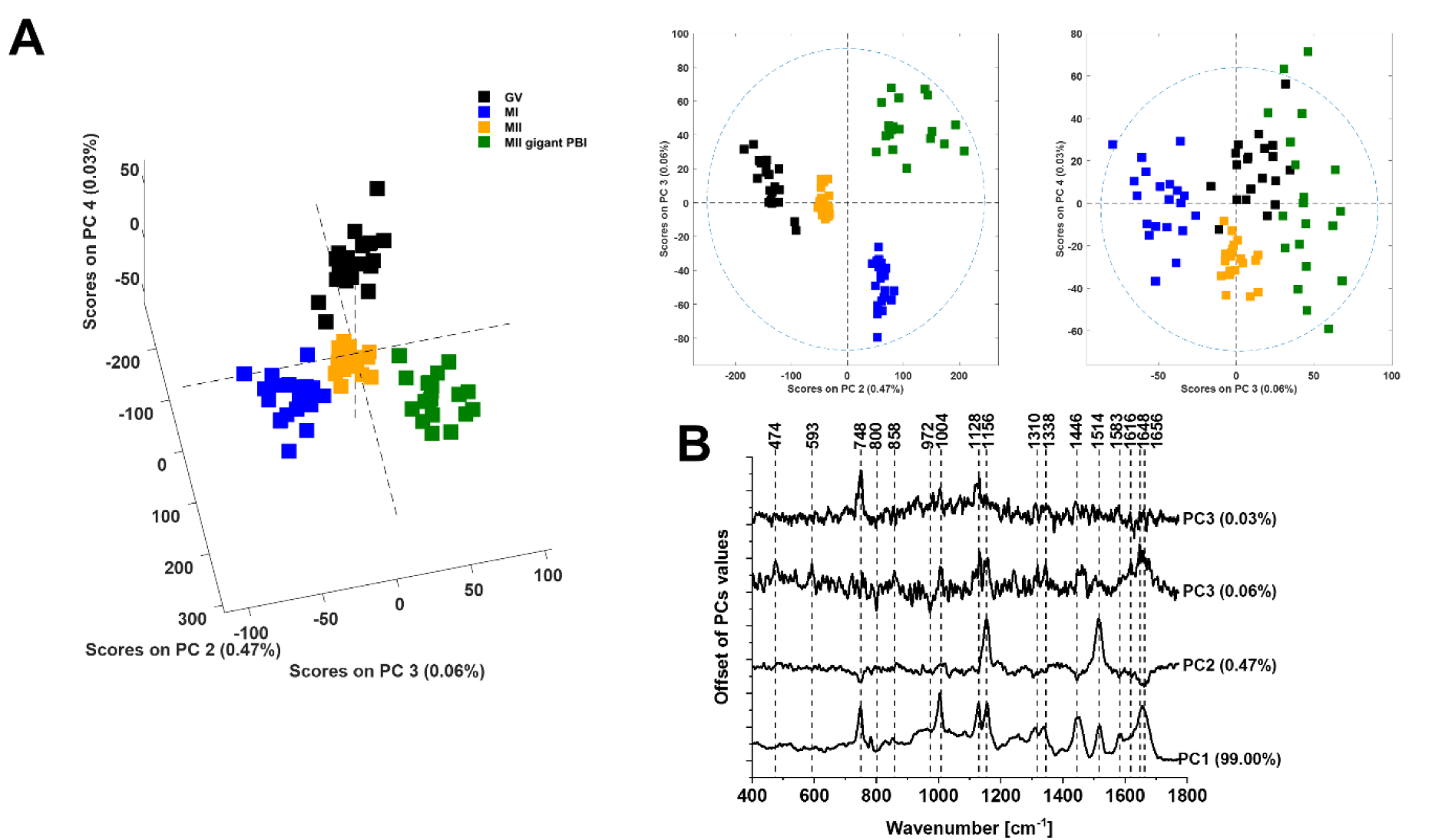
Principal components analysis of Raman data for different stages of maturation GV, MI, MII, MII giant PB. The scatter plot of the score values of Raman spectra for the first, second and third principal components (PCs) from the GV (black squares), MI (blue squares), MII (yellow squares), MII with giant PBI (green squares) of human oocytes (A). The loadings plots of PC1, PC2 and PC3 (B). Data was acquired at 532 nm with 0.3 second integration time. The 20 average Raman spectra were recorded across a line of each oocyte. The total number of 80 average spectra from the human oocyte cells were analyzed. Preprocessing PCA mode: SNV normalization.

## Conclusions

We investigated the biochemical composition of specific organelles in human oocytes at various stages of maturation (GV, immature MI, MII with the first polar body, MII with giant polar body and vacuoles) using Raman imaging. Our study demonstrated that label-free Raman imaging can provide detailed biochemical characterization and functional status of structures such as the zona pellucida, perivitelline space, polar body, vacuoles, and organelles like mitochondria, cytoplasm, and the nucleus in oocyte cells.

Through the integration of Raman imaging and chemometric classification via Cluster Analysis, we conducted a comprehensive biochemical analysis of the proteomic, lipidomic, and DNA profiles of the oocytes. Our findings highlight that cytochrome *c* is a crucial protein in oocyte mitochondria, essential for both respiration and apoptosis. Normal functioning of oocyte cells requires cytochrome *c* in its redox-balanced forms: reduced and oxidized. We showed that the oxidized form of cytochrome *c* predominates in oocyte mitochondria, playing a vital role in the proper electron transport chain. This allows effective electron shuttling between complex III, cytochrome *c*, and complex IV, facilitating controlled oxidative phosphorylation and ATP production.

Moreover, our study suggests that Raman imaging can be utilized for rapid fertility testing by assessing the mitochondrial status in oocytes, potentially providing a valuable tool for reproductive medicine.

## Materials and Methods

### Ethical approval

Written informed consent was obtained from the oocyte donors attending the Infertility Clinic Salve Medica, Łódź, Poland. The study was approved by the Bioethical Committee of the Medical University in Lodz, Poland (No RNN/83/23/KE).

### Human oocytes

Human oocyte samples were obtained from 6 donors. Women who fulfilled the following inclusion criteria were eligible: age between 26 and 32 years, body mass index (BMI) between 18 and 30 kg/m², and basal FSH less than 10 IU/L (cycle day 2– 5). All participants were administered 150–225 IU recombinant FSH (Gonal-f, Merck Serono, Germany) from the third day of the menstrual cycle. Follicles were monitored through transvaginal sonography 5–6 days after gonadotropin stimulation. When the follicular diameter reached 13–14 mm, Cetrotide (0.25 mg/d; Merck Serono, Germany) was prescribed. When three dominant follicles reached >17 mm, 250 µg (6,500 IU) of hCG (Ovitrelle, Merck Serono, Germany) was administered for the final maturation of oocytes and ovulation. Oocyte retrieval was performed 36 hours after the trigger. The removal of cumulus cells in a process called oocyte denudation was performed with Hyaluronidase (Gynemed Germany). Unfertilized abnormal/immature cells (3 GV cells, 3 MI cells, and 1 abnormal cell with a large polar body) as well as unfertilized normal oocytes(2 MII), which were designated for disposal because more were collected than planned for fertilization, were designated for the study on confocal Raman micro-spectroscopy analysis.

### Confocal Raman micro-spectroscopy

The nine human native live oocytes from six donors were put between two CaF_2_ Raman grade window slides with a 150 μm micron spacer (Harrick, MSP-150-M25) in G-MOPS™ PLUS (Vitrolife, Goteborg, Sweden) to perform Raman evaluation.

The confocal Raman microscope (WITec (alpha 300 RSA+), Ulm, Germany) in the Laboratory of Laser Molecular Spectroscopy, Lodz University of Technology, Poland was applied to record Raman spectra and images. The excitation laser at 532 nm was focused on the sample to the laser spot of 1 µm and was coupled to the microscope via an optical fiber with a diameter of 50 µm. The average laser excitation power was 10 mW, and the collection time was 0.3 second (with EMCCD gain) for Raman images. Raman images were recorded with a spatial resolution of 1 × 1 µm. A typical Raman map of an oocyte cell consists of 22500 Raman spectra (map size 150 x 150 μm). The Raman spectrometer was calibrated every day prior to the measurements using a silica plate with a maximum peak at 520.7 cm^-1^.

### Data analysis

The obtained Raman data were analyzed by Cluster Analysis using the Project Plus (WITec GmbH, Germany), Origin 2018 (Origin Lab, USA) and Principal Component Analysis (PCA) analysis was performed using MATLAB (MathWorks, USA) with PLS-Toolbox (Eigenvector Research Inc., USA). PCA analysis is a dimension reduction analysis that allows the identification of patterns in high dimensional data, expressing the data in such a way as to highlight their similarities and differences.

## Author Contributions

Conceptualization: H.A. J.M.S.; Investigation: J.M.S.; Methodology: J.M.S., H.A., K.M.; Sample preparation: J.M.S., K.M.; Funding acquisition: H.A.; Writing – original draft: H.A., J.M.S.; Manuscript reviewing: H.A., J.M.S., K.M., J.S-H., R.W-J, B.S.; Supervision: J.S-H.; H.A. B.S. All authors have read and agreed to the published version of the manuscript.

## Funding

This work was supported by the National Science Centre of Poland (Narodowe Centrum Nauki, UMO-2021/43/B/ST4/01547) and Medical University of Lodz grant no 503/1-089-03/503-11-002.

## Data Availability Statement

The raw data underlying the results presented in the study are available from Lodz University of Technology Institutional Data Access. Request for access to those data should be addressed to the corresponding author.

## Conflicts of Interest

The authors declare no competing interest. The funders had no role in the design of the study; in the collection, analyses, or interpretation of data; in the writing of the manuscript, or in the decision to publish the results.

## Notes

### Competing Interest Statement

The authors have declared no competing interest.

